# Ovarian cancer cell cluster inhibits pancreatic cancer progression

**DOI:** 10.1101/2025.01.26.634970

**Authors:** Maodi Liang, Rui Fan, Cuizhe Wang, Kejian Gong, Xiao Lin, Mengyuan Zhao, Huizi Zhang, Lili Xu, Qin Liu, Yurui Su, Chunmei Cui, Xingkai Liu, Jun Zhang, Qinghua Cui

## Abstract

Pancreatic cancer is the most lethal cancer and all of the current cancer treatment strategies fail at this disease. It is thus urgently important to explore novel therapeutics. We previously revealed that it can be significantly inhibited by ovarian cancer cell by both *in vitro* and *in vivo* experiments. It was reported that cancer cell cluster can greatly enhance the efficiency in metastasis model compared to single cancer cell. Given this observation, here we proposed that ovarian cancer cell cluster could inhibit pancreatic cancer more efficiently than single ovarian cancer cell. As a result, animal experiments demonstrated that the injection of both low and high doses of SKOV3 ovarian cancer cluster show significantly higher efficiency in inhibiting pancreatic cancer xenografts than single SKOV3 ovarian cancer cell. Moreover, single cell RNA-sequencing analysis revealed that ovarian cancer cell cluster treatment inhibited pancreatic cancer by promoting inflammation response by both activating the NF-κB pathway in pancreatic cancer cells and substantially shifting macrophages in tumor microenvironments (TME) to M1 phenotype. In summary, this study proposed and validated a novel potential therapeutic for pancreatic cancer.

## Introduction

Pancreatic cancer is one of the most lethal human malignancies with increasing incidence and mortality rates in many countries^1^. In United States, it was reported that pancreatic cancer has become the third leading cause of cancer-related death after surpassing breast cancer, and is projected to become the second one when overtaking colorectal cancer before 2040^2^ or by 2030^3^. To date, although the overall survival of patients with pancreatic cancer has been improved by best available systemic therapies including surgery, chemotherapy, radiation therapy, interventional therapy and immunotherapy, it still remains poor, with a 5-year survival rate of only ∼11% after diagnosis due to therapeutic failures^4^ as most patients present with advanced disease^5^. During the last decade, there has been some novel paradigms in pancreatic cancer therapeutics. Next-generation sequencing and bioinformatics analysis have revealed some driver mutations and aberrant pathways in pancreatic cancer, which further identified novel targets and promoted the development of targeted therapy^6,7^, however, which only do benefit to a small fraction of patients and improve limited survival time. For example, only 25% of the patients host somatic/germ line mutations in DNA Damage Repair (DDR) pathway^6^. For immunotherapies such as ipilimumab (cytotoxic T-cell antigen-4 inhibitor), PD-L1 monoclonal antibody (immune checkpoint inhibitor) and CAR-engineered T cells (CAR-T cells), they often show good effects on melanoma^8,9^ and haematological malignancies^10^, but traditionally did not lead to response in pancreatic cancer due to immunosuppressive tumor microenvironment (TME)^6^. Therefore, heating the “cold” immune system of pancreatic cancer would activate its response to immunotherapies^6^. For example, an animal experiment showed that tetanus toxin derived from injected listeria bacteria can induce memory T cells and then activate cytotoxic T cells in TME^11^. Although new techniques in pancreatic cancer early detection, subtyping, management and treatment have improved its survival, pancreatic cancer still represents one of the most lethal cancers with increasing incidence and mortality rate. Its treatment is still a huge challenge and novel treatment approaches are required.

Bioinformatics is one new technique which play roles in various biological and medical fields^12^. Using reverse-transcriptomics based bioinformatics technique, we previously found that the gene expression signature of pancreatic cancer is significantly anti-correlated with that of ovarian cancer, suggesting ovarian cancer cell could have ability of defensing pancreatic cancer. We then confirmed this hypothesis using *in vitro* and *in vivo* experiments and proposed a novel strategy for pancreatic cancer treatment, combating pancreatic cancer with ovarian cancer cell (CCC or C^3^) therapeutics^13^. Meanwhile, Chen et al. reported direct tumor killing and antitumor immunity by cancer cell-based vaccine^14^, also suggesting a possibility of combating cancer using cancer cells.

Recently, we have learned that “*Experimentally aggregating tumor cells into clusters induced a >15-fold increase in colony formation ex vivo and a >100-fold increase in metastasis formation in vivo*” from a study for polyclonal breast cancer metastases^15^. Based on this observation, here we proposed that ovarian cancer cell cluster would greatly further enhance the efficiency in defensing pancreatic cancer compared to single ovarian cancer cell in the C^3^ therapeutics. As a result, we confirmed this hypothesis. The injection of both low and high doses of SKOV3 ovarian cancer cluster show significantly higher efficiency in inhibiting pancreatic cancer xenografts than single SKOV3 ovarian cancer cell. Moreover, single cell RNA-sequencing analysis revealed that ovarian cancer cell cluster treatment can both activate the inflammatory activity on tumor cells and substantially alternate tumor microenvironments (TME) to inhibit pancreatic cancer. In summary, this study proposed and validated a novel potential therapeutic for pancreatic cancer (**c**ombating pancreatic **c**ancer with ovarian **c**ancer cell **c**luster, C^4^).

## Results

### SKOV3 ovarian cancer cell cluster greatly enhanced the inhibition of pancreatic tumor development than single SKOV3 ovarian cancer cell

Cluster organization originates from multicellular grouping of oligoclonal tumors, improving the survival of tumor cell colonization and subsequently increasing metastatic events^15-17^. Combined with our previous results that only a small number of ovarian cancer cells are distributed in pancreatic tumor tissues after two weeks of delivering single SKOV3 cells^13^, here we hypothesize that enhancing the colonization ability of SKOV3 by aggregating it into clusters may potentially improve CCC therapeutics. To assess the therapeutic efficacy of multicellular seeding on pancreatic cancer, we processed pancreatic cancer xenografts with ovarian tumor cell (OTC) clusters (**Figure** 1a). Specifically, OTCs were obtained by isolating and digesting the SKOV3 xenografts from nude mice. After incubation for 24 hours in ultra-low adhesion dishes, OTCs were allowed to aggregate to form OTC clusters (**Figure** 1b). Intratumoral injection of single OTCs and OTC clusters was performed on pancreatic cancer every two days, with two concentrations of low dose (LD, 1×10^5^) and high dose (HD, 3×10^5^), respectively. Gemcitabine, a first-line drug for pancreatic cancer therapy, was used as a positive control to evaluate the proposed new therapeutics. The results showed that after two weeks of treatment with OTC, whether single OTCs or OTC clusters, effectively inhibited the growth of pancreatic cancer and reduced the tumor weight (**Figure** 1c-f), which is consistent with our previously observations^13^. Notably, at the same cell density, OTC clusters seeding significantly suppressed the growth of pancreatic cancer compared to the single OTCs treatment group (**Figure** 1c). After two weeks of OTC clusters delivery, tumor volume of pancreatic cancer were reduced by 27.4% in the low-dose group (648.86 ± 77.50 mm^3^ vs. 894.12 ±109.00 mm^3^, *p* = 0.003) and by 28.6% in the high-dose group (609.02 ± 72.51 mm^3^ vs. 853.20 ±66.85 mm^3^, *p* < 0.001) (**Figure** 1c-e). Meanwhile, tumor weight were also significantly reduced in both the low-dose (0.60 ± 0.11 g vs. 0.79 ±0.08 g, *p* = 0.014) and high-dose cluster groups (0.53 ± 0.10 g vs. 0.76 ±0.08 g, *p* = 0.005) compared with the single OTCs controls (**Figure** 1f). These data indicate that OCT cluster exerts an enhanced anti-pancreatic cancer effect. Additionally, we did not observe any alterations in the weight of nude mice during the OTC intervention (**Figure** 1g). In summary, our results demonstrate that the C^4^ therapeutics (combating pancreatic cancer with OTC cluster) shows a more potent effect in combating pancreatic cancer than the C^3^ therapeutics (combating pancreatic cancer with single OTCs) we proposed previously.

**Figure 1.**
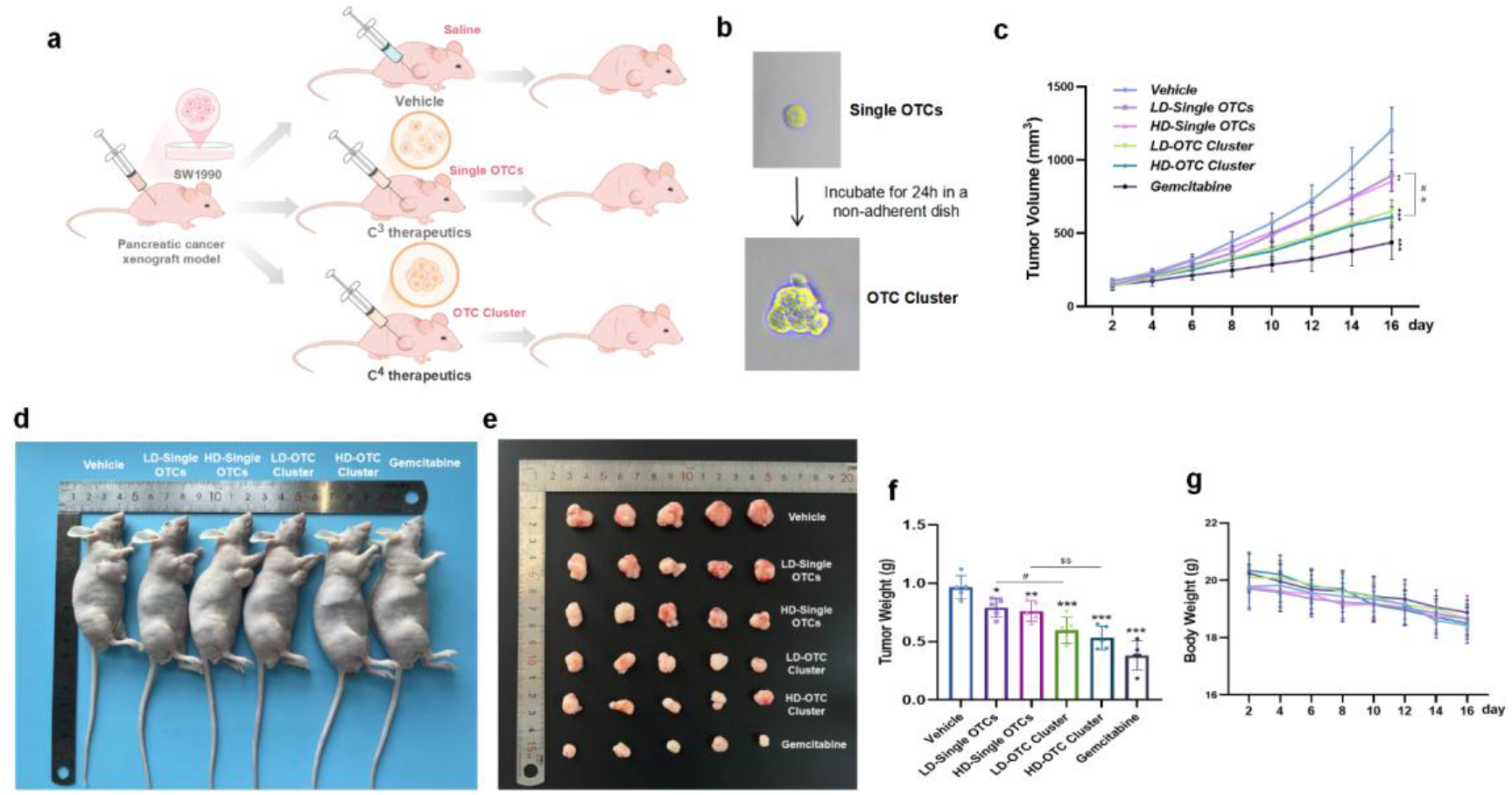
OTC cluster enhanced the inhibition of pancreatic tumor development than single OTCs. **a**, Experimental design. SW1990 cells were used to establish the pancreatic cancer subcutaneous xenograft model, and single OTC (ovarian tumor cell) and OTC cluster were treated separately, with saline as a control. **b**, Formation of OTC cluster (200×). OTCs were isolated from ovarian cancer xenografts and aggregated in ultra-low adsorption culture plates for 24 h. **c**, Pancreatic tumors in nude mice after intervention. Intratumoral injection of single OTCs and OTC clusters every two days, with low dose and high dose. **d**, Isolation of pancreatic tumors. **e**, Tumor growth curve during intervention. The points and bars represent means ± SD. **f**, Tumor weights are shown as means ± SD. **g**, Body weight changes of mice during intervention. n =5. LD, low dose. HD, high dose. OTC, ovarian tumor cell. \**p* < 0.05, \*\**p* < 0.01, ^##^*p*< 0.01, ^$$^*p*< 0.01. *, compared to the vehicle group.

### C4 therapeutic inhibits the pancreatic tumor through activating inflammatory

To elucidate the mechanisms of the proposed C4 therapeutic strategy, we performed single-cell RNA sequencing (scRNA-seq) on tumor tissues from three experimental groups: a vehicle control group, a group receiving single OTC treatment, and a group receiving OTC cluster treatment. Our analysis revealed substantial alterations in the pancreatic tumor cell compartment (**Figure** 2a). As expected, we observed a reduction in the proportion of pancreatic tumor cells following both single OTC and OTC cluster treatments (**Figure** 2d). Specifically, the single OTC treatment and the OTC cluster treatment inhibited 56.0% and 81.3% of pancreatic tumor cells when being compared to the vehicle group (**Figure** 2d). Moreover, clustering analysis revealed four distinct cell subpopulations in pancreatic tumor cells (**Figure** 2b, e). Notably, the relative abundance of these subpopulations varied across treatment groups (Figure 2c), indicating differential therapeutic effects. For instance, IL2RG+ subpopulation was markedly enriched in the OTC cluster treatment group (**Figure** 2c). Subsequent pathway enrichment analysis of marker genes characterizing IL2RG+ subpopulation revealed significant activation of inflammatory pathways, including the NF-κB and IL-17 signaling pathways (**Figure** 2f). These findings suggest that the observed inhibition of pancreatic tumor development by the C4 therapeutic may be mediated, through the activation of inflammatory pathways within the tumor cells.

**Figure 2.**
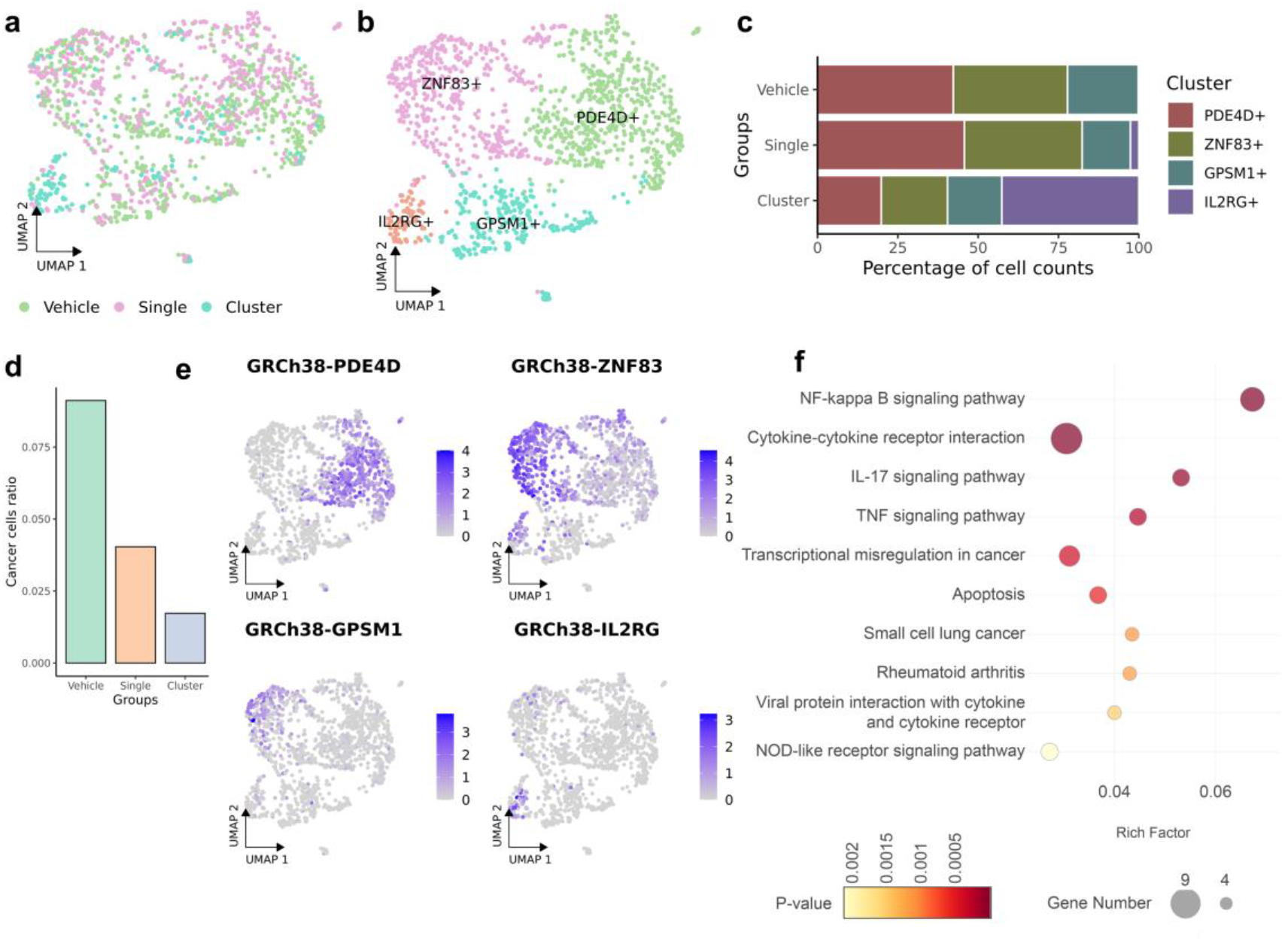
Single-cell RNA sequencing analysis of pancreatic tumor cells. **a**, UMAP visualization of total pancreatic tumor cells colored by the three experimental groups. **b**, identification of subpopulations of pancreatic tumor cells. **c**, proportional abundance of subpopulations in the three experimental groups. **d**, proportional representation of pancreatic tumor cells within the experimental groups. **e**, expression of subpopulation specific marker genes. **f**, pathway enrichment analysis (KEGG) of the subpopulation 3 specific genes.

### Tumor microenvironment contributes to the C4 therapeutic of pancreatic tumor

The tumor microenvironment (TME), a complex and dynamic ecosystem, plays a critical role in cancer development, progression, and response to therapy ^18^. Thus, we next investigated the TME alterations induced by the C3 and C4 therapeutics. Using the scRNA-seq data and the clustering algorithm, we identified eight distinct cell subpopulations within TME (**Figure** 3a-b). These subpopulations were characterized based on their transcriptomic profiles and the expression of cell type-specific marker genes, including neutrophils, macrophages, T cells, B cells, fibroblasts, smooth muscle cells (SMCs), endothelial cells, and epithelial cells (**Figure** 3b, d). Notably, the relative proportions of these cell types varied across the experimental groups, with macrophages and fibroblasts exhibiting most pronounced changes (**Figure** 3c).

**Figure 3.**
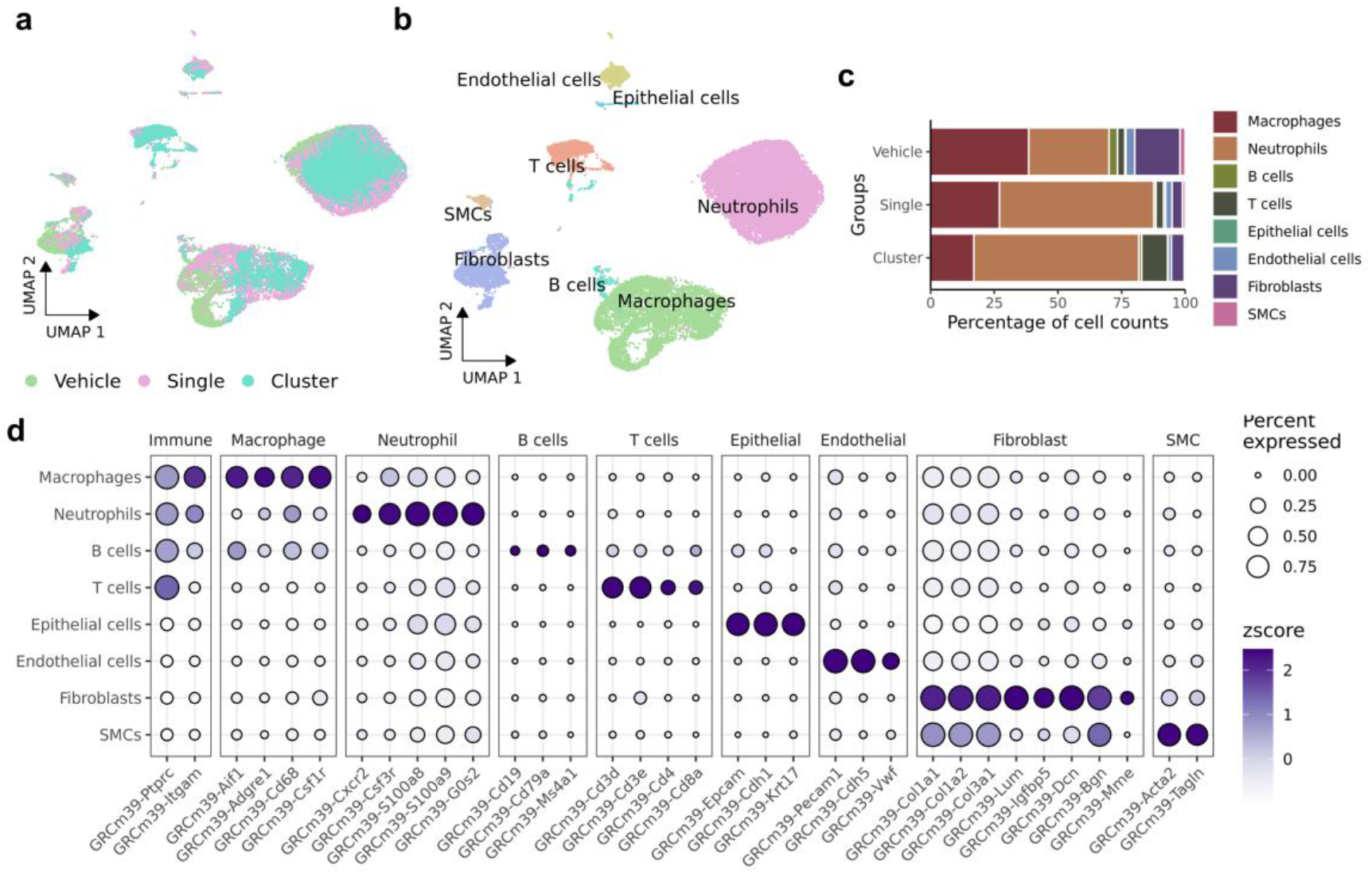
Single-cell RNA sequencing analysis of tumor microenvironment (TME). **a**, UMAP visualization of TME cells colored by the experimental groups. **b**, identification of major cell types within TME. **c**, proportional abundance of TME cell types across the 3 experimental groups. **d**, expression patterns of the cell type-specific marker genes.

Macrophages, key components of the TME, are recognized for their diverse and often paradoxical roles in cancer development and progression, influencing both tumor growth and anti-tumor immunity ^19^. To further understand the heterogeneity of macrophages in our model, we identified four distinct macrophage subpopulations, whose relative abundance varied across the experimental groups (**Figure** 4a-c). Analysis of marker gene expression within these subpopulations revealed several genes of interest (**Figure** 4d). For example, *Nos2* encodes for inducible nitric oxide synthase (iNOS), which is a key enzyme in the production of nitric oxide (NO) (**Figure** 4d). Numerous studies have demonstrated the significant influence of NO on cancer development and treatment ^20^. Furthermore, we observed that *Folr2* has recently been identified as a marker for a tissue-resident macrophage (TRM) population and interacts with tumor-infiltrating CD8+ T cells ^21^ (**Figure** 4d). Furthermore, we assessed the M1 and M2 polarization signatures of individual macrophages using a scoring system derived from established gene sets ^22,23^ (**Figure** 4e). As a result, both the single OTC treatment and the cluster OTC treatment have higher M1 score and M2 score than the vehicle treatment. Moreover, the cluster OTC treatment have higher M1 score than the single OTC treatment. Our analysis revealed that both the single treatment and the cluster treatment induced a shift towards an M1-like polarization state in macrophages. In addition, we observed a decrease in both phagocytosis and angiogenesis scores in the treatment groups ^24^ (**Figure** 4f), suggesting a potential reduction in macrophage-mediated tumor-promoting activities.

**Figure 4.**
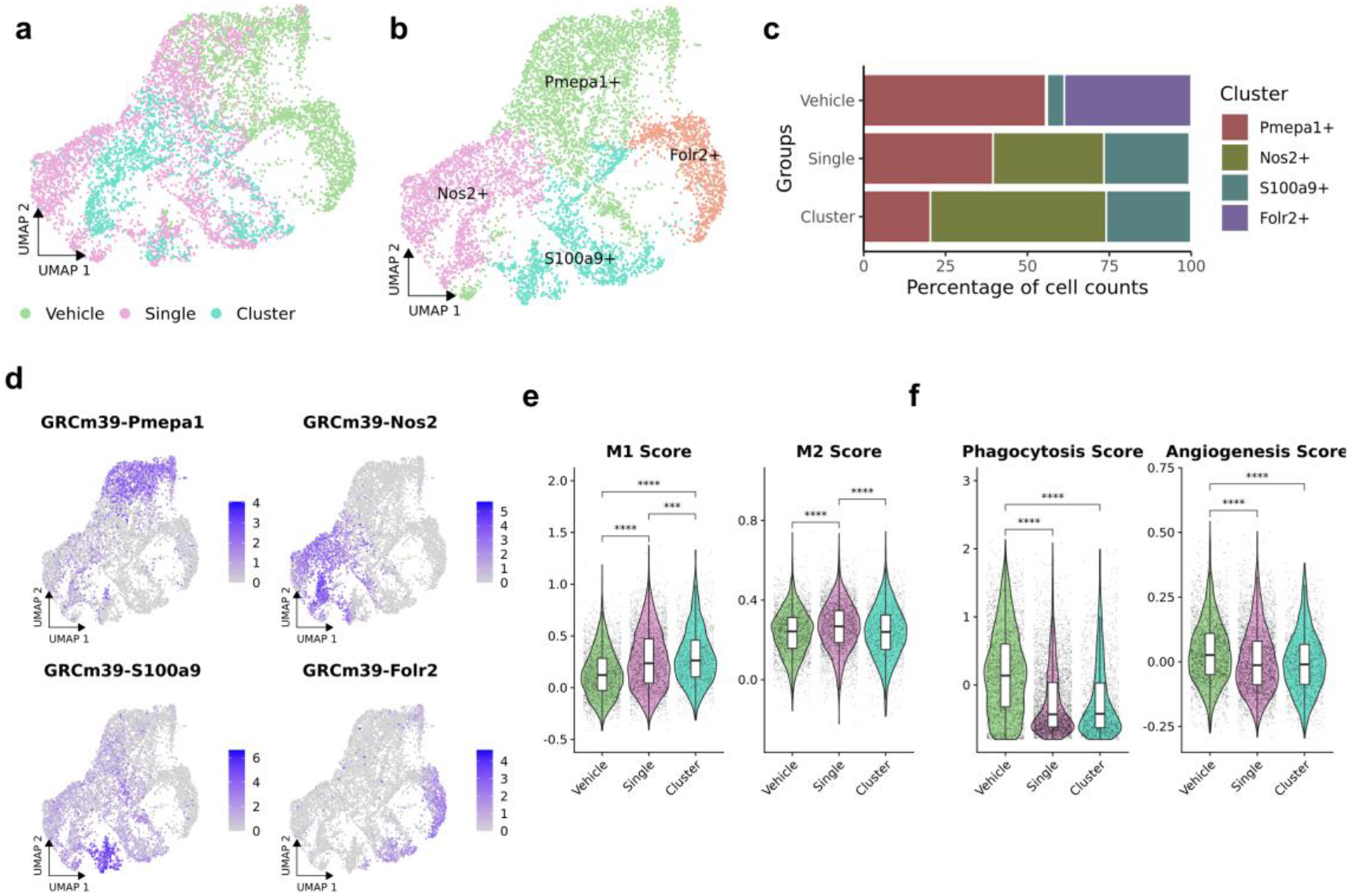
Single-cell analysis of macrophage subpopulations. **a**, UMAP visualization of macrophages colored by the experimental groups. **b**, identification of macrophages subtypes. **c**, proportional abundance of different cell types across the experimental groups. **d**, expressions pattern of the subpopulation marker genes. **e**, distributions of M1 score and M2 score in different groups. **f**, distributions of phagocytosis score and angiogenesis score in different groups. The Wilcoxon rank test were employed on e and f. ****, *p*<0.0001. ***, *p*<0.001.

To further investigate the stromal compartment of the TME, we performed single-cell RNA sequencing analysis of fibroblasts, revealing substantial heterogeneity within this cell population (**Figure** 5a). Using an unbiased clustering algorithm, we identified distinct fibroblast subtypes (**Figure** 5b), and characterize them based on their myofibroblast (myCAF), inflammatory fibroblast (iCAF), and antigen-presenting fibroblast (apCAF) signatures using established scoring systems ^23^ (**Figure** 5d). As a result, the fibroblast exhibited differential representation across the experimental groups (**Figure** 5c), suggesting that the C3 (the single OTC treatment) and C4 therapeutics (the cluster OTC treatment) may modulate fibroblast activation and phenotypic polarization. Specifically, we observed an enrichment of Lrrc4+ fibroblasts and Trem3+ fibroblasts in C3 and C4 treatment groups. These findings suggest that the C3 and C4 therapeutics may not only directly target tumor cells but also indirectly influence the TME by modulating the functional landscape of fibroblasts

**Figure 5.**
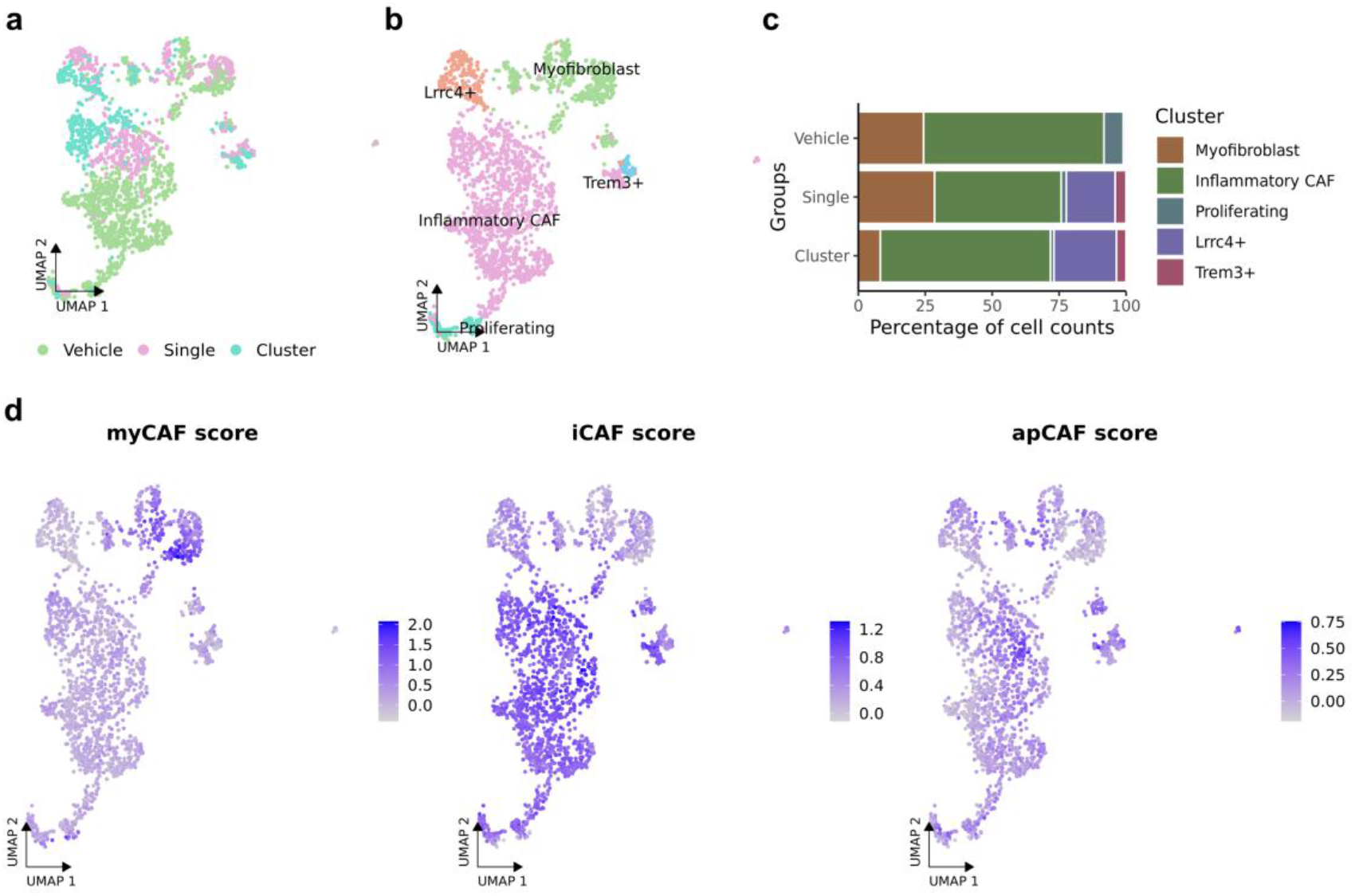
Single-cell analysis of fibroblast heterogeneity. **a**, UMAP visualization of fibroblasts colored by the experimental groups. **b**, identification of fibroblast subtypes. **c**, proportion of different cell types in different groups. **d**, distributions of myCAF score, iCAF score and apCAF in different groups.

## Discussion

Pancreatic cancer is one of the most lethal cancers and a major cause of cancer death in the world^25^. Current treatment approaches usually fails at pancreatic cancer, although there are some advances in its therapy such as targeted therapies^26^, T cell activating^27^, radiopharmaceuticals^28^, and mRNA vaccine^29^ etc. Therefore, novel therapeutics is emergently needed. Thanks for the rapid accumulation of omics data, we previously predicted that the gene signature of pancreatic cancer is anti-correlated with that of ovarian cancer. Based on this observation, we proposed and then confirmed by in vitro and in vivo experiments that ovarian cancer cell can defense pancreatic cancer, and then proposed a novel therapeutic, (**c**ombating pancreatic **c**ancer with ovarian **c**ancer cells, C^3^) therapeutics. It was reported that cancer cell cluster can greatly enhance the efficiency in metastasis model compared to single cancer cell. Based on this observation, this study proposed and confirmed by animal experiments an updated C^3^ therapeutics, combating pancreatic cancer with ovarian cancer cell cluster, that is the C^4^ therapeutics. The C^4^ therapeutics could be a potential effective approach for pancreatic cancer treatment. Moreover, scRNA-seq analysis revealed that the C^4^ therapeutics reduced the pancreatic cancer cells more efficiently, which may be mediated by enhancing inflammation response through activating NF-κB and IL-17 signaling pathways. It is well known that NF-κB is a critical factor in promoting inflammation and causing cancer development and progression^30^. The scRNA-seq analysis here suggest the other side of the coin, activating inflammation in established tumor may inhibit cancer progression. Besides pancreatic cancer cells, scRNA-seq analysis also shows that both treatments have significant effects on TME cells. For example, both treatments enhanced the M1 score of TME macrophages and the cluster OTC treatment enhanced more significantly, which also suggest that promoting inflammation in established tumor through shifting macrophages to M1 phenotype may inhibit cancer progression.

Although significances have been found, there are still a number of important issues should be addressed in the future. Firstly, the efficacy of the C^4^ therapeutics should be validated by more preclinical models as different models may address unique challenges such as progression, metastasis, and stromal heterogeneity^31^. For example, pancreatic cancer patients with bone metastasis have an extremely low overall survival, and it is quite difficult to treat bone metastasis^32^. Therefore, it is necessary to investigate whether the C^4^ therapeutics still works in this situation by animal models of bone metastasis of pancreatic cancer. Secondly, currently the mechanisms of the C^3^ and C^4^ therapeutics are not clear and should be explored, which is critical for the applications of this therapeutics on human patients. Some new omics techniques such as single-cell sequencing^33^ and spatial transcriptome^34^ may be of great helps for exploring the mechanism of the proposed therapeutics. In addition, it also remains unclear whether this therapeutic still works on other types of cancer, that is, it is important to investigate the generalization of this therapeutics.

## Methods & Materials

### Animals

4-6-week male BALB/c nude mice were obtained from SPF biotechnology Co. Ltd. (Beijing, China), and housed at the Laboratory Animal Center of Shihezi University after a week of adaptive feeding. Mice were fed *ad libitum* during the experimental intervention. All animal protocols were approved by the Medical Ethics Committee of the First Affiliated Hospital of Shihezi University School of Medicine and complied with ethical regulations related to animal studies.

### Cell Lines

Human pancreatic cancer cell line SW1990 was purchased from ATCC Co., Ltd (USA). Lentiviral vector-transfected GFP-tagged SKOV3 cells were performed as we previously described^13^. SW1990 cells and SKOV3-GFP cells were cultured in DMEM/F12 (Gbico, USA) containing 10 % fetal bovine serum (FBS, Gbico, USA) and 1 % Penicillin-Streptomycin (Gbico, USA). Cells were cultured at 37°C in a humidified incubator containing 5 % CO_2_. 80 % cell confluence was used to establish the subcutaneous graft model.

### Isolation of OTCs and cluster incubation

SKOV3-GFP cells (8×10^6^/100 μL per mice) in logarithmic growth phase were subcutaneously inoculated into the right axilla of nude mice. Subsequently, the ovarian cancer tumor tissue was aseptically dissected and placed in DMEM/F12 medium containing 10 % FBS and 2 % penicillin-streptomycin at 4 °C. The tumor tissue was minced into pieces approximately 1-2 mm^3^ in size using surgical instruments. In DMEM/F12 medium containing type IV collagenase (1.5 mg/mL, Solarbio, China), digestion was carried out at 37°C for 1 hour. After filtration, the dissociated OTCs were collected by centrifugation. For cluster incubation, transfer the isolated OTCs into ultra-low adsorption 96-well plates (Labselect, China) and allow them to aggregate for 24 hours. Collect the OTC clusters into tubes, and suspend with saline for experiment after a brief centrifugation.

### Establishment of pancreatic cancer xenograft model

SW1990 cell suspension (5×10^6^/100 μL per mice) of logarithmic growth phase in the total volume of 0.1 mL was inoculated subcutaneously into the right posterior axillary of nude mice. Treatment was started when the tumor volume reached about 150mm^3^. The tumor volume was calculated with a standard formula: width^2^×length×0.52. After completion of the intervention, the xenografts were weighted, excised, fixed in formalin or frozen at -80°C for further experiments.

### Single-cell RNA-sequencing and analysis

Single-cell RNA-sequencing (scRNA-seq) is performed by Beijing Boao Crystal Code Biotechnology Co., Ltd., China. In brief, fresh pancreatic cancer tissues are separated after the experiment. Using the 10× Genomic platform, single-cell gel beads are generated in the emulsion. The prepared single-cell suspension is loaded into the 10× Genomic single-cell capture system to construct a cDNA library with 10× tags. Short-read NGS sequencing is performed on an Illumina sequencer. The data obtained can obtain the gene expression profile and difference analysis at the single-cell level through Cell Ranger.

Analysis of scRNA-seq data was performed using the Seurat package (version 5.0.2) in R (version 4.3.3). Initial quality control involved filtering cells based on the proportion of mitochondrial gene transcripts to remove low-quality cells. Subsequently, data normalization was conducted, followed by dimensionality reduction using the standard Seurat pipeline, which includes principal component analysis (PCA). The Uniform Manifold Approximation and Projection (UMAP) algorithm was then employed for non-linear dimensionality reduction and visualization of single cells in a two-dimensional space. Gene set enrichment analysis was performed using the Weighted Enrichment Analysis Tool (WEAT) algorithm ^35^.

### Quantification and Statistical Analysis

All data was expressed as the mean ± SD. R (version 4.2) was used for all statistical analyses. For a comparison between two groups, the T-test was used. For comparisons between three or more groups, the one-way ANOVA (one-tailed) was used.

## Credit authorship contribution statement

QC proposed the original idea and JZ designed the experiment. ML performed the experiments. CW and KG performed part of the experiments. XL, HZ, MZ, LLX, QL, YS, and CC provided helps in animal experiments or data analysis. QC and ML wrote the draft manuscript. QC, JZ, and XL supervised the study.

## Acknowledgments

We thank Prof. Lixiang Xue for providing the SKOV3 cell line.

